# Thermal stress induces positive phenotypic and molecular feedback loops in zebrafish embryos

**DOI:** 10.1101/2021.04.07.438623

**Authors:** Lauric Feugere, Victoria F. Scott, Quentin Rodriguez-Barucg, Pedro Beltran-Alvarez, Katharina C. Wollenberg Valero

## Abstract

Aquatic organisms must cope with both rising and rapidly changing temperatures. These environmental changes can affect numerous traits, from molecular to ecological scales. Biotic stressors can induce the release of chemical cues which trigger behavioural responses in other individuals. In this study, we infer whether abiotic stressors, such as fluctuating temperature, may similarly propagate stress responses between individuals in fish not directly exposed to the stressor. To test this hypothesis, zebrafish (*Danio rerio*) embryos were exposed for 24 hours to fluctuating thermal stress, to medium in which another embryo was thermally stressed before (“stress medium”), and to a combination of these. Growth, behaviour, and expression of a panel of genes were used to characterise the thermal stress response and its propagation between embryos. Both high temperatures and stress medium significantly accelerated development and altered embryonic behaviour. Thermal stress significantly decreased the expression of the antioxidant gene SOD1, eight hours after the end of exposure. Of note, we found that the expression of sulfide:quinone oxidoreductase (SQOR), likewise a part of the antioxidant metabolism relevant in vertebrate stress response, and of interleukin-1β (IL-1β), involved in the immune response, were significantly altered by stress medium. This study illustrates the existence of positive thermal stress feedback loops in zebrafish embryos that induce stress in conspecifics. This evidence that thermal stress due to fluctuating, high temperatures can be propagated may be relevant for species found in high densities, either in aquaculture or in the natural environment, in a context of global change.

## Introduction

Temperature is the abiotic ‘ecological master factor’ regulating the biology of ectotherms (Brett, 1971; López-Olmeda & Sánchez-Vázquez, 2011). Ectotherms face thermal cycles that shape their biological rhythms by modulating survival, growth, and by triggering irreversible changes on thermal tolerance ranges of adults (Colinet et al., 2015; Kingsolver et al., 2015; Lim et al., 2017; López-Olmeda & Sánchez-Vázquez, 2011; Schaefer & Ryan, 2006). Those natural, regular thermal fluctuations experienced during early development are necessary to maximise fitness-relevant traits and thermal tolerance (Hall & Warner, 2020b; Kingsolver et al., 2015; Lim et al., 2017; Schaefer & Ryan, 2006). Besides higher average temperatures, global warming will be characterised by both greater thermal variability and more extreme thermal events of longer duration and magnitude (Pörtner et al., 2019; Vasseur et al., 2014). These altered thermal rhythms may pose higher risks to species than simply higher mean temperatures, due to the asymmetry of the thermal fitness curve (Colinet et al., 2015; Saxon et al., 2018; Vasseur et al., 2014). Here, we exposed zebrafish embryos to several peaks of higher temperatures that consisted in thirteen exposures to +5°C within 24 hours. Although this thermal regime departs from natural, environmental relevant extreme heatwaves, it serves to model the effects of repeated thermal stress upon the stress response of zebrafish embryos.

Fish as ectothermic vertebrates are susceptible to changes in the thermal environment, particularly to higher temperatures close to their upper thermal limits (Araújo et al., 2013; I. J. Morgan et al., 2001; Paaijmans et al., 2013). Early developmental stages have narrower thermal ranges than adults (Skjærven et al., 2014). Temperature regimes during development have irreversible effects as they modulate subsequent stages, making early fish embryos vulnerable or “bottleneck” stages in the context of climate change (Pörtner & Peck, 2010; Scott & Johnston, 2012; Villamizar et al., 2012). Examples for persistent effects of temperature changes during development may involve alterations of swimming performance and cardiac anatomy (Dimitriadi et al., 2018), masculinisation (Ribas et al., 2017) and increased mortality (Hosseini et al., 2019), which can shape the future trajectory of populations. Altogether, this illustrates a rising concern about the response of fish to global change, particularly at early stages of development.

The zebrafish (*Danio rerio*) is now emerging as a model organism to study the effects of thermal stress, including at the molecular level (Brown et al., 2015; Clark et al., 2011; Long et al., 2012; Luu et al., 2020). Adult zebrafish are eurythermal, naturally tolerating a wide range of temperatures (16.5-38.6°C) with optimal temperature around 27-28.5°C. They may face natural thermal variations of ∼5°C daily, and from 6 to 38°C seasonally (Engeszer et al., 2007; López-Olmeda & Sánchez-Vázquez, 2011; Spence et al., 2008). However, early stages of zebrafish only tolerate minimum temperatures of 22-23°C and maximum temperatures of around 32°C to develop normally (Pype et al., 2015; Schirone & Gross, 1968; Schnurr et al., 2014). Warm-adapted species such as the zebrafish that are living near their upper thermal limit may be among the ‘losers’ of climate change (Somero, 2010). Of note, the thermal biology of zebrafish is conserved in laboratory populations, in spite of laboratory domestication, which makes them an adequate model organism to investigate the effects of climate change in the laboratory (Brown et al., 2015; R. Morgan et al., 2019).

Besides the above-mentioned effects on sex ratio, direct mortality and adult phenotype, responses to thermal stress at the early stages of development involve altered behaviour, developmental rate, and altered gene expression related to the physiological stress response. Behavioural thermoregulation is one major thermoregulatory process in ectotherms (López-Olmeda & Sánchez-Vázquez, 2011). This is evident from a tightly controlled thermotaxis response in response to heat in zebrafish larvae (Haesemeyer et al., 2015, 2018). Heat stressed zebrafish larvae display more anxiety-like behaviours (Bai et al., 2016) and transient hyperactivity (Yokogawa et al., 2014). High temperatures also accelerate the embryonic development in zebrafish embryos as shown by the 2.8-fold increase in somitogenesis frequency across a 20-30°C range or the twice faster development at 33°C compared to 26°C (Hallare et al., 2005; Long et al., 2012; Schröter et al., 2008). Repeated exposure to sublethal temperatures, however, may depress development (Hall & Warner, 2020a). At the molecular level, heat stress leads to numerous molecular effects from the accumulation of reactive oxygen species (ROS) (Madeira et al., 2016; Vinagre et al., 2012) to changes in global gene expression patterns (Logan & Buckley, 2015; Long et al., 2012; Ribas et al., 2017) along the hypothalamic-pituitary-interrenal (HPI) axis (Alsop & Vijayan, 2009). For example, Cu/Zn-superoxide dismutase I (SOD1) neutralises oxygen radicals to protect cells from oxidative stress (Cheng et al., 2018; Wang et al., 2016), and is heat inducible in zebrafish embryos (Icoglu Aksakal & Ciltas, 2018). Another less well studied enzyme, sulfide:quinone oxidoreductase (SQOR) is upregulated by heat stress in half-smooth tongue sole (*Cynoglossus semilaevis*) (Guo et al., 2016) and by hypoxia in Nile tilapia (*Oreochromis niloticus*) (J. H. Xia et al., 2018). SQOR expression is co-induced by both cold stress and hypoxia in zebrafish embryos (Long et al., 2012, 2015). The innate immune system is challenged by both high temperature and thermal fluctuations in teleost early stages (Mariana & Badr, 2019; Zhang et al., 2018). For instance, interleukin-1β (IL-1β) is upregulated in zebrafish raised at high temperature but experiencing an immune challenge at low temperature (Zhang et al., 2018) and in zebrafish embryos exposed to high temperature (Icoglu Aksakal & Ciltas, 2018). This gene plays a central role in the stress response, having neuromodulatory and behavioural functions (Goshen & Yirmiya, 2009; Metz et al., 2006; Vitkovic et al., 2000).

Largely overlooked to date, however, is the question of whether the response to thermal stress can be transmitted to other individuals. Fish use chemical communication to alert others of a threat using so-called ‘alarm cues’ (released after skin damage) or ‘disturbance cues’ (released without injury following a biotic stressor) (Jordão & Volpato, 2000). Exposure to conspecific predation-related disturbance cues induces stress-like responses in several fish species (Barcellos et al., 2011, 2014; Bett et al., 2016; Ferrari et al., 2008; Jordão & Volpato, 2000; Toa et al., 2004), including zebrafish (Barcellos et al., 2014). Importantly, even fish early stages are capable of such chemical communication as they respond to alarm cues (Atherton & McCormick, 2015, 2017; Oulton et al., 2013; Poisson et al., 2017). Despite the wealth of information on biotic stress-induced stress propagation, this phenomenon is only known from a few abiotic stressors, such as low pH, acute fasting or handling (Abreu et al., 2016; Feugere et al., in review), but in response to heat is only known from crayfish (Hazlett, 1985). Stress induces the release of metabolites into the environment, including hormones of the HPI axis (Barcellos et al., 2014; McGlashan et al., 2012), CO_2_ (McGlashan et al., 2012), respiratory byproducts, catecholamines, or nitrogenous metabolic products such as urea (Bairos-Novak et al., 2017; Giaquinto & Hoffmann, 2012; Henderson et al., 2017; Hubbard et al., 2003), but less is known about their effects on communication. Distinct from alarm and disturbance cues, we introduced the term ‘stress metabolite’ referring to such cues induced without injury as a byproduct of the response to abiotic stress (Feugere et al., in review).

In this contribution we infer whether thermal stress can be propagated to naive receivers. To test this hypothesis, zebrafish embryos were exposed to independent and combined treatments of thermal stress and medium putatively containing stress metabolites induced by this thermal stress. Zebrafish embryo behaviour, growth rate, and expression of genes involved in the immune response (IL-1β) and antioxidant pathways (SOD1, SQOR) were investigated. We expected that (i) fluctuating high temperatures, similar to constant higher temperatures, trigger developmental, behavioural and molecular stress responses in zebrafish embryos. Second (ii), we hypothesised that these responses could induce a positive feedback loop in naive receiver embryos, which are however (iii) not elicited by non-stress metabolites. It should be emphasised that our work primarily aimed to investigate the existence of stress propagation induced by repeated thermal stress, in a context of growing concern for a future more stressful environment, rather than study the effects of realistic heatwaves.

## Materials and Methods

### Experimental design

For detailed zebrafish husbandry and breeding methods, see Supplementary Information. Just before the beginning of experimental treatments, several zebrafish embryos per clutch were photographed to estimate the median initial stage. Exposure began around the blastula stage (median stage: 2.75 hpf, at 512-cells). Fertilised healthy embryos (with chorion) were selected and individually placed into 0.2 mL 8-strip PCR tubes prefilled with 200 µL of 1X E3 medium.

In a two-factorial design, embryos were exposed to different combinations of the two factors, temperature stress and temperature-induced stress medium. For this purpose, embryos were exposed to either thermal stress or control temperature protocols within a thermocycler, in either fresh medium or medium where embryos had previously been exposed to thermal stress and containing putative stress metabolites. All experimental treatments are detailed in Figure 1 and Table S1. The thermal stress protocol spanned 16.25 hrs, divided into thirteen 75-min series of temperature fluctuations between 27, 29, 32, 29, and 27°C, with each temperature step being maintained for 15 min. Thermal stress mimicked +5°C temperature peaks over zebrafish optimal temperature (a total of n = 13 peaks) reaching the sublethal temperature of 32°C (Scott & Johnston, 2012). A recovery time of 7.75 hrs at 27°C followed the fluctuating temperatures period to reach a total incubation time of 24 hrs. The control thermal protocol was a steady 27°C for 24 hrs. E3 media following control or thermal stress conditions were reused for ‘control medium’ and ‘stress medium’ treatments, respectively. In total this yielded five treatments: control (C, control protocol with fresh medium), control medium (CM, control protocol with reused medium from C), thermal stress (TS, thermal stress protocol with fresh medium), stress medium (SM, control protocol with reused medium from TS), and thermal stress + stress medium (TS+SM, thermal stress protocol with reused medium from TS). Used media were immediately re-used for treatments containing putative stress or control metabolites, respectively. As development in zebrafish accelerates with higher temperature (Kimmel et al., 1995) but decelerates with darkness (Bucking et al., 2013; Villamizar et al., 2014), additional control embryos were monitored after longer incubation times: 31 hrs (C31), 37 hrs (C37), and 46 hrs (C46). These times were adjusted for darkness-raised embryos to reach the stages of prim-6 (25 hpf), prim-16 (31 hpf), and late pharyngula (35 - 42 hpf), respectively. Initial, final, and total exposure times were used to standardise each procedure. After incubation, transparent embryos were deemed alive and kept for subsequent steps. Before and after each exposure, embryo media were sampled to measure pH and O_2_ saturation levels that could impact embryos in the used media. Oxygen levels were averaged from 1-2 min measurements using a glass vial equipped with an oxygen sensor spot (OXSP5, sensor code: SC7-538-193, Pyroscience GmbH) and connected to the FireSting O_2_ Fiber Optic Oxygen Meter (FSO2-4, Pyroscience GmbH) and Oxygen Logger software (Pyroscience GmbH). Constant medium pHs were verified using the Fisherbrand(tm) accumet(tm) AB150 pH Benchtop Meters.

**Figure 1.**
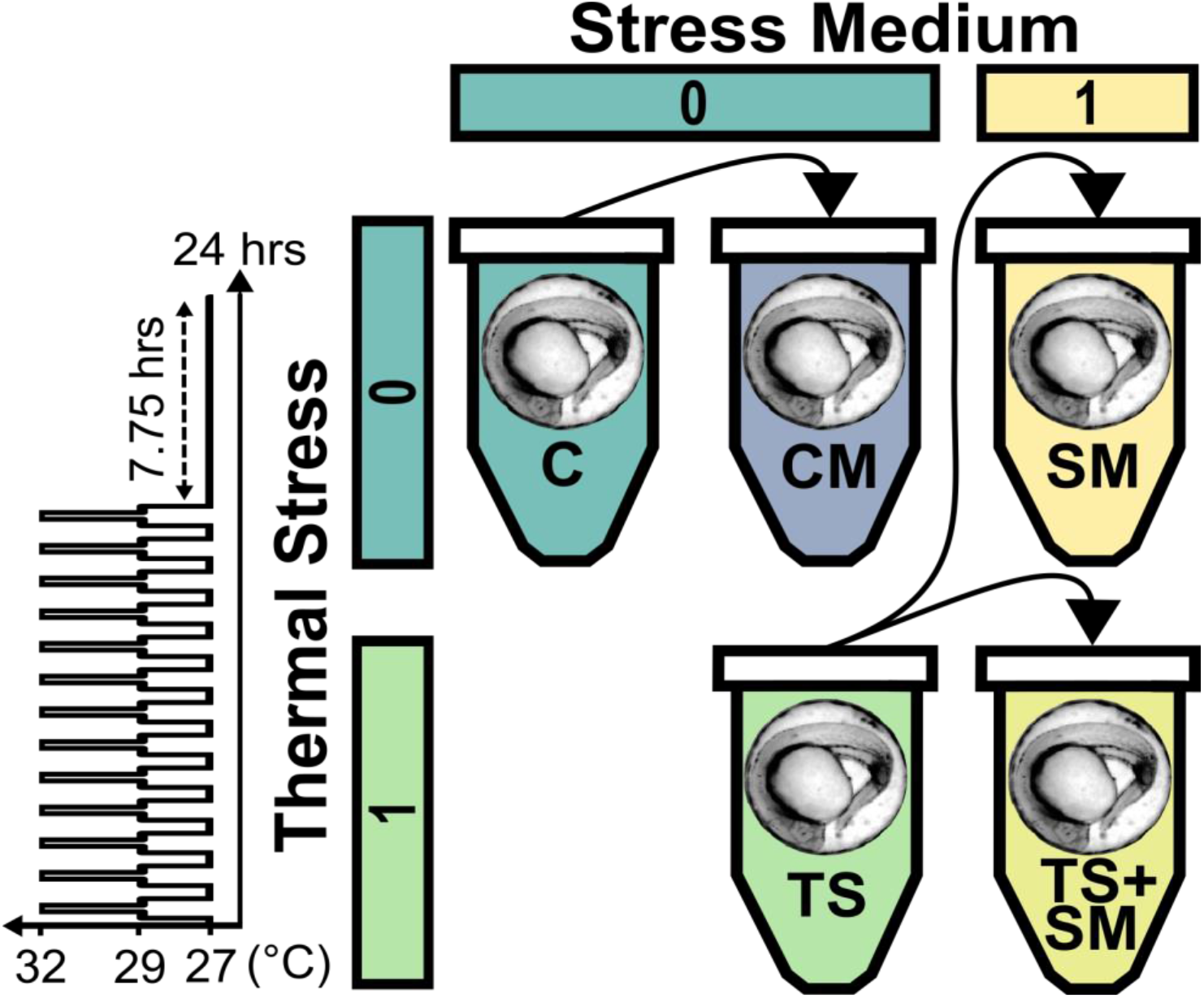
Overview of experimental design. In a two-way factorial design (represented by 0 and 1 for factor levels, respectively), zebrafish embryos (*Danio rerio*) were exposed to either fluctuating thermal stress (inset diagram) followed by a recovery period (dashed arrow line) or constant control temperature (27°C), in combination with fresh E3 medium or stress medium containing putative stress metabolites produced by one previously thermally-stressed embryo. An additional treatment was incubated at 27°C in a control medium containing metabolites from an embryo previously exposed to control conditions. Plain black arrows indicate medium transfer in which a new embryo was incubated. CM: control medium, C: control, SM: stress medium, TS: thermal stress, TS+SM: thermal stress + stress medium.

### Analysis of phenotypic data

Phenotypic analyses were conducted for all eight treatments. Embryos were placed in a watch glass vial and videoed by small batches using the camera setup (see Supplementary Information) over 15-30 seconds. Embryos were placed under light (similar intensity across measurements) to elicit a startle-like response after exposure in darkness. Behavioural data was analysed using Danioscope (Noldus). When possible, several videos were recorded for each embryo clutch and behavioural measurements were averaged for each individual embryo. Analysis of behavioural responses were conducted on the video-length standardised burst activity percentage (percentage of the time – from the total measurement duration – the embryo was moving). Final embryonic stages were estimated following Kimmel et al. (1995) from several photographs of embryos within their chorions using the criteria somite number, yolk extension to yolk ball ratio, presence and morphology of otoliths, tail aspect, presence of lens primordium, presence and position of the cerebellum relatively to the eyes, and pigmentation. The growth index was calculated as in equation 1.

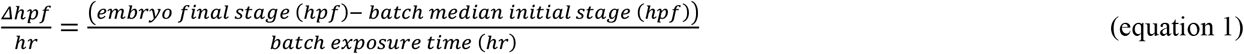

Statistical analyses were conducted in RStudio (RStudio Team, 2020). Outliers among behavioural values were excluded from statistical analyses using Tukey’s method with a 1.5 interquartile range cut-off. First, the effects of stress medium and thermal stress predictors across the C, TS, SM, and TS+SM treatments were evaluated. Shapiro-Wilk and Levene’s tests were used to evaluate normality and homogeneity of variances, respectively. Normalisation methods were compared using the *BestNormalize* R package (Peterson & Cavanaugh, 2020). The growth index was parameterised using an order normalising transformation (Shapiro-Wilk’s P = 0.98, Levene’s P = 0.27) and analysed by a two-way ANOVA with thermal stress and stress medium as predictors, followed by *post-hoc* pairwise Student’s t-tests between control C and each experimental condition. The burst activity percentage and final stage (in hpf) could not be normalized and were analysed for the effects of both predictors using nonparametric Scheirer-Ray-Hare tests from the *rcompanion* R package (Mangiafico, 2018), followed by pairwise *post-hoc* Wilcoxon-Mann-Whitney tests. The final embryonic developmental period was coded as *segmentation* or *pharyngula* and analysed for the effects of thermal stress, stress medium as factors and initial stages as covariate using a generalised logistic model followed by pairwise *post-hoc* comparisons using the *emmeans* R package (Lenth, 2019).

Additional pairwise comparisons were performed of all response variables of CM against C and SM. To determine whether behavioural effects of treatment were related to developmental acceleration, pairwise comparisons of behaviour of control embryos (C) were compared to that of older control embryos (C31, C37, C46) using Kruskal-Wallis tests. Fourth, burst activity percentages of embryos from stress medium and thermal stress treatments were compared against that of control embryos incubated for longer times using pairwise Wilcoxon-Mann-Whitney tests. Multiple comparisons were corrected using Bonferroni adjustments. Cohen’s |d| were obtained from the distribution used to compute the statistical analyses and calculated using the *effsize* r package (Torchiano, 2016) or according to Lenhard & Lenhard (2016), respectively. Effect size was qualified based on thresholds given in Sawilowsky (2009): very small: |d| > 0.01, small: |d| > 0.2,medium: |d| > 0.5, large: |d| > 0.8, very large: |d| > 1.20, huge: |d| > 2.0.

### Gene expression

Gene expression analyses were conducted for CM, C, SM, TS, and TS+SM treatments (n = 3 pooled biological replicates of 60 embryos per treatment). Embryos were snap-frozen at -80°C immediately after experimental treatments. Total RNA was extracted using a High Pure RNA isolation kit (Sigma-Aldrich) following the manufacturer’s recommendations. cDNA was synthesized using Superscript II™ Reverse Transcriptase (Invitrogen, Life Technologies Ltd.) with sample randomisation. TaqMan® Gene Expression Assays (ThermoFisher Scientific) and 2X qPCR Bio Probe Hi-ROX (PCRBiosystems) were used to quantify the expression of three genes of interest (SQOR, SOD1, and IL-1β) normalised to two reference genes (ef1-α, β-actin). The effects of stress medium and thermal stress on the Log_2_ 2^-ΔΔCT^ (Log_2_ fold-change) values were investigated using the *eBayes* and *lmFit* functions within the *limma* package (Ritchie et al., 2015) within the *Bioconductor v*.*3*.*11* (Ihaka & Gentleman, 1996) R environment. Next, pairwise *post-hoc* comparisons on C *versus* SM, TS, or TS+SM, and CM *versus* C or CM were performed using pairwise moderated t-tests with Bonferroni adjustments. Effect sizes (Cohen’s |d|) were calculated as above. More details on the gene expression analysis are given as Supplementary Information.

## Results

### Phenotypic effects of thermal stress and its propagation

First, the phenotypic effects of fluctuating thermal stress and of stress medium treatments were analysed. Embryonic growth indices were significantly accelerated by stress medium (small effect size, F = 6.291, P = 0.0128), thermal stress (large effect size, F = 75.502, P < 0.0001), and their combination (very large effect size, F = 7.498, P = 0.0067, Figure 2a, Table 1). *Post-hoc* tests revealed that TS (t = -7.9874, P < 0.0001), SM (t = - 3.6784, P = 0.0012), and the combined treatment TS+SM (t = -7.2413, P < 0.0001) all accelerated growth, compared to the control C (Table S2). The growth acceleration was accompanied by a median advancement in embryonic stages of 3 to 9 hours, resulting in a switch from the segmentation to the pharyngula stage, compared to controls C and CM (see Supplementary Information, Figure S1a-b, Tables S3-S4). Treatments had no obvious effect on mortality.

**Table 1.** Effects of thermal stress and stress medium on the growth index of pre-hatching zebrafish embryos. A two-way ANOVA was used to test the effects of the predictors (thermal stress and stress medium) on changes in growth index. Effect size is computed as Cohen’s |d| and interpreted according to thresholds given in Sawilowsky (Sawilowsky, 2009). Effect sizes of significant p-values (P ≤ 0.05) are shown in bold.

| **Model terms** | **F statistic** | **P** | **$\vert d\vert $** | **Effect size** |
| --- | --- | --- | --- | --- |
| Stress Medium | 6.291 | 0.0128 | 0.29 | small |
| Thermal Stress | 75.502 | 0.0001 | 1.14 | large |
| Thermal Stress x Stress Medium | 7.498 | 0.0067 | 1.27 | very large |

**Figure 2.**
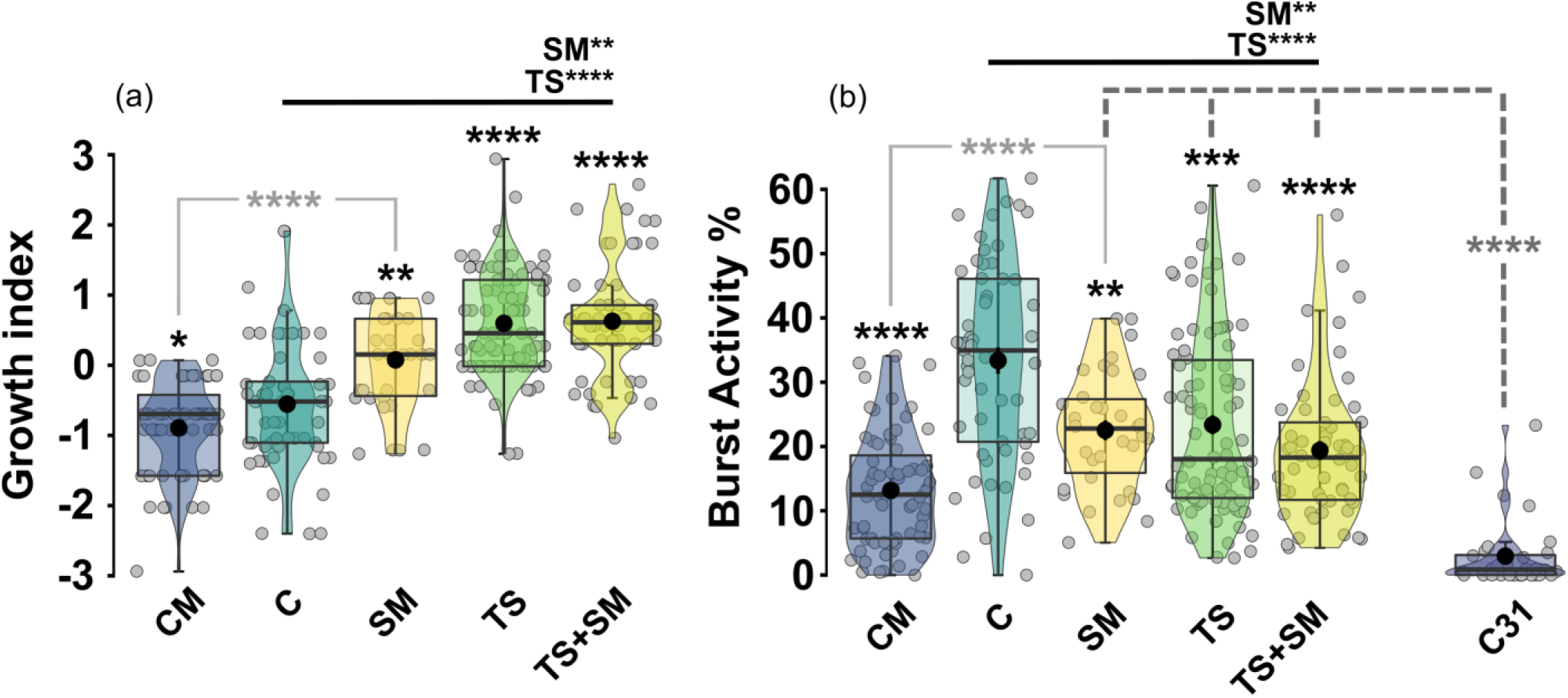
Thermal stress and stress medium accelerate the growth index (a) and lower the burst activity percentage (b) of pre-hatching zebrafish embryos. CM: control medium (*n* = 67), C: control (*n* = 57), SM: stress medium (*n* = 34), TS: thermal stress (*n* = 90), TS+SM: thermal stress + stress medium (*n* = 56), C31: control after 31 hrs of incubation (reaching prim-6 stage). Boxes represent median, 25%-75% quartiles, and whiskers are minimum and maximum values within 1.5 IQR (interquartile range). Grey dots represent individual data points and black dots represent mean values. Effects of predictors (thermal stress and stress medium) were tested with two-factorial tests. Significant predictors are indicated in bold above horizontal black lines with *: *P* ≤ 0.05, **: *P* ≤ 0.01, ***: *P* ≤ 0.001, ****: *P* ≤ 0.0001. Significance of pairwise post-hoc tests is likewise indicated in asterisks, for treatments against C (bold black), between SM and CM (light grey), or between over-developing treatments (SM, TS, TS+SM) and C31 (dashed bar with dark grey asterisk).

We reasoned that any observed effects of stress medium may be due to regularly excreted metabolites, which accumulate towards the end of each treatment independently of thermal stress and would be up-concentrated in SM and TS+SM treatments. This warranted the use of an additional control, the ‘control medium’ (CM), that is, medium that had previously hosted control embryos. This control helped us exclude oxygen saturation and pH as confounding effects, which were also independently measured at the beginning of treatments and in all cases fell within zebrafish natural tolerance ranges (Strecker et al., 2011; Zahangir et al., 2015). We therefore assessed whether stress medium, that is, medium where embryos had been exposed to thermal stress, evoked different effects compared to control medium. Embryonic growth in CM was compared to growth in C and SM. Embryos in CM grew slightly slower than control embryos (t = 2.3472, P = 0.0418) and much slower than embryos in SM (t = -6.7215, P < 0.0001, Figure 2a, Table S2).

Next, we investigated the behavioural startle-like response to light (Figure 2b). Burst activity percentages were significantly lowered by both predictors stress medium (H = 9.3222, P = 0.0023) and thermal stress (H = 17.008, P < 0.0001), whereas the interaction term was not significant (H = 1.8193, P = 0.1774, Table 2). *Post-hoc* comparisons showed that embryos treated with SM (W = 1,387, P = 0.0018), TS (W = 3,548, P = 0.0003), and TS+SM (W = 2,455, P < 0.0001) all displayed lower burst percentages compared to control C. Embryos exposed to CM showed even stronger decline in burst activity percentages compared to C (W = 3,287, P < 0.0001) and SM (W = 1,745, P < 0.0001, Table S5).

**Table 2.**
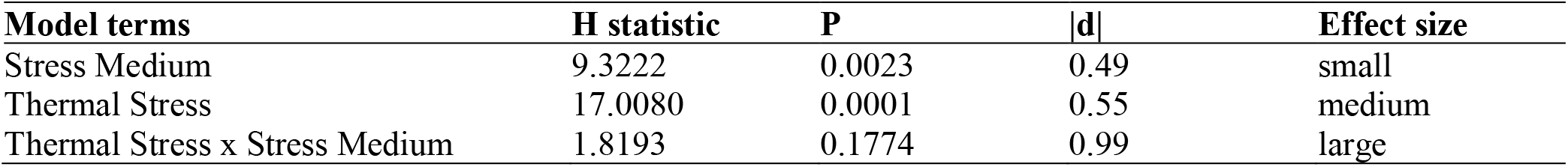
Effects of thermal stress and stress medium on the burst activity percentage of pre-hatching zebrafish embryos. A two-way Scheirer-Ray-Hare test was used to test the effects of the predictors (thermal stress and stress medium) on burst activity percentages across treatments. Effect size is computed as Cohen’s |d| and interpreted according to thresholds given in Sawilowsky (Sawilowsky 2009). Effect sizes of significant p-values (P ≤ 0.05) are shown in bold.

| Model terms | H statistic | P | \|d\| | Effect size |
| --- | --- | --- | --- | --- |
| Stress Medium | 9.3222 | 0.0023 | 0.49 | small |
| Thermal Stress | 17.0080 | 0.0001 | 0.55 | medium |
| Thermal Stress x Stress Medium | 1.8193 | 0.1774 | 0.99 | large |

Considering the aforementioned thermal stress-induced growth acceleration, stage-dependent effects of behaviour were investigated in control embryos incubated for 24, 31, 37, or 46 hrs. We observed that burst activity percentages significantly decreased with development in pre-hatching control embryos raised for 31, 37, and 46 hrs compared to those incubated for 24 hrs (P < 0.0001, Figure S2c, Table S6). However, neither final stages (P = 0.1627) nor growth index (P = 0.5027) were correlated with burst activity percentages across treated embryos (Table S5). Moreover, pairwise tests showed that stressed embryos under development accelerated by TS, SM, and TS+SM treatments were significantly more active than control embryos reaching the same median stage of prim-6 in C31 (P < 0.0001, Figure 2b, Table S7).

### Effects of thermal stress and its propagation on stress response-related gene expression

The whole-embryo expression of three stress-inducible candidate genes (IL-1β, SOD1 and SQOR) was analysed (Figure 3, Tables 3 and S8). Thermal stress was not a significant predictor for the expression of either IL-1β or SQOR, but SOD1 expression was lower in the TS treatment compared to the control C (very large effect size but marginal significance, t = 2.76, P = 0.045). In contrast, stress medium significantly reduced IL-1β expression (very large effect size, t = 2.28, P = 0.038) and increased SQOR expression levels (huge effect size, t = -3.54, P = 0.003), but had no effects on SOD1 expression.

**Table 3.** Effect of thermal stress and stress medium on gene expression of pre-hatching zebrafish embryos. Effects of model terms (thermal stress and stress medium) on the expression of IL-1β, SOD1 and SQOR obtained via moderated t-tests (t-statistic) with *lmfit* and *eBayes* in the *limma* R package. B describes the log-odds of gene expression. Effect size is computed as Cohen’s |d| and interpreted according to thresholds given in Sawilowsky (Sawilowsky 2009). Effect sizes of significant p-values (P ≤ 0.05) are shown in bold.

| Model terms | t | B | P | \|d\| | Effect size |
| --- | --- | --- | --- | --- | --- |
| **IL-1β** |  |  |  |  |  |
| Thermal Stress | 1.03 | -4.70 | 0.310 | 0.49 | medium |
| Stress Medium | 2.28 | -3.95 | 0.038 | 1.2 | very large |
| **SOD1** |  |  |  |  |  |
| Thermal Stress | 1.65 | -4.27 | 0.108 | 1.08 | large |
| Stress Medium | 0.32 | -6.16 | 0.757 | 0.17 | small |
| **SQOR** |  |  |  |  |  |
| Thermal Stress | -0.17 | -4.99 | 0.869 | 0.11 | small |
| Stress Medium | -3.54 | -1.67 | 0.003 | 2.43 | huge |

**Figure 3.**
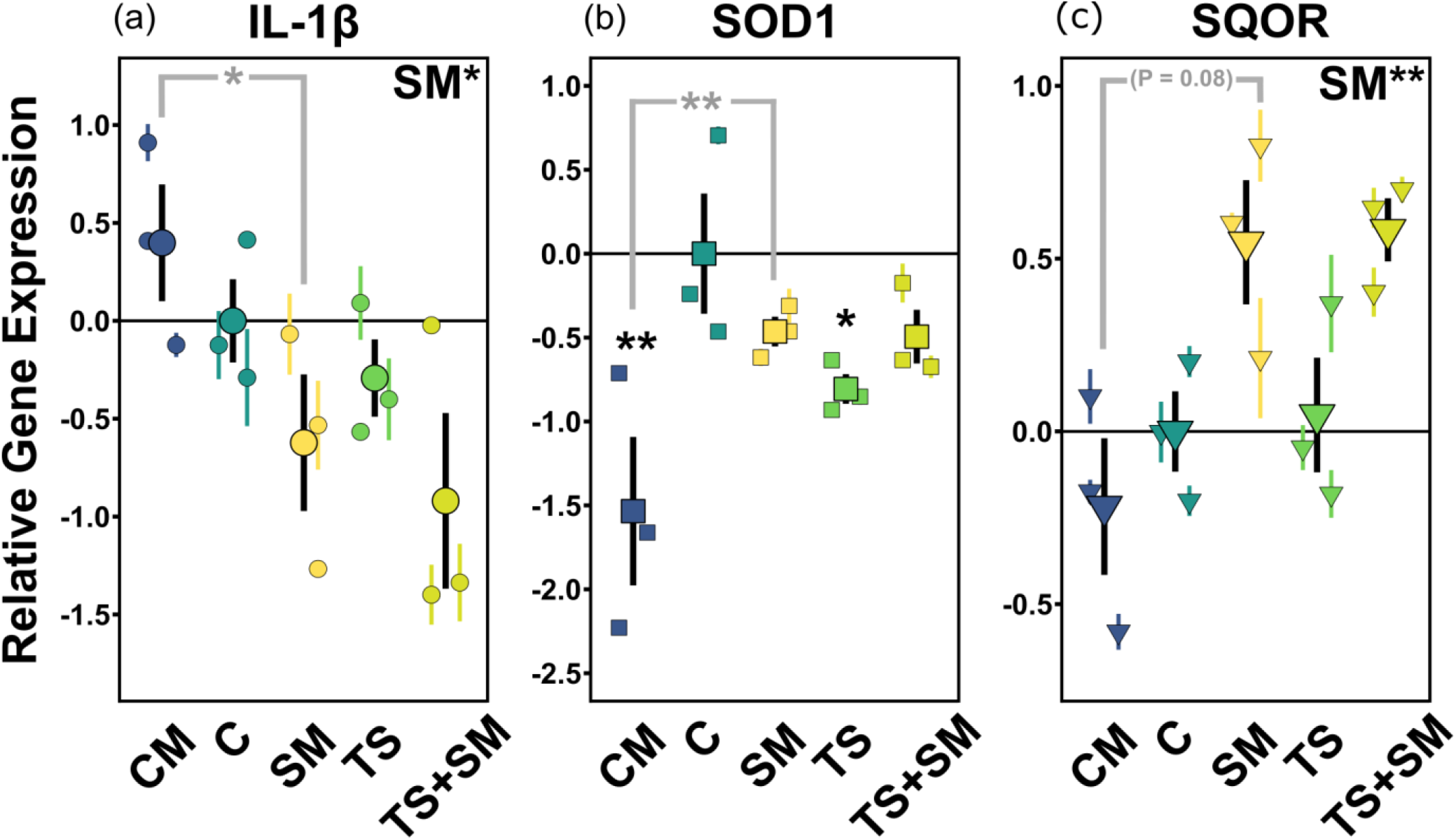
Stress medium alters stress-related gene expression of zebrafish embryos. (a) IL-1β (interleukin-1β), involved in immune response. (b) SOD1 (superoxide dismutase 1), involved in antioxidant response. (c) SQOR (sulfide:quinone oxidoreductase), involved in metabolism and antioxidant response. Jittered symbols represent mean Log_2_ 2^-ΔΔCT^ ± SE (coloured bars) values of each biological replicate. Black symbols represent mean Log_2_ 2^-ΔΔCT^ ± SE (black bars) values of each biological treatment. Expression is relative to control C (mean values as black horizontal lines). CM: control medium, C: control, SM: stress medium, TS: thermal stress, TS+SM: thermal stress + stress medium. Annotations on top right corners represent the factorial effects of thermal stress (TS) and stress medium (SM) on C, SM, TS, TS+SM. Black asterisks above each mean value only show the pairwise comparisons against the control C. Comparisons between CM and SM are represented in grey by asterisks and dashed horizontal lines. Statistics were computed using moderated t-tests (Limma, Bioconductor, R) with Bonferroni adjustment with *: P ≤ 0.05, **: P ≤ 0.01. Sample size: *n* = 3 biological replicates for each treatment, with 60 pooled embryos per each replicate.

Pairwise tests revealed that SOD1 gene expression in control medium (CM) was much lower compared to C (huge effect size, t = -3.78, P = 0.003), but did not differ from C for the expression of SQOR and IL-1β. Second, pairwise tests showed that stress medium treatment (SM) triggered different gene expression responses compared to CM in all three studied markers. SM significantly reduced the expression of IL-1β (very large effect size, t = 2.62, P = 0.034) and increased that of SOD1 (very large effect size, t = -2.64, P = 0.034), whereas there was a only a non-significant trend towards increased SQOR expression (P = 0.08). Relative to the control, the gene expressions of IL1-β and SQOR in SM showed opposite patterns compared to CM. Taken together, our gene expression results suggest that incubation in stress medium leads to reduced IL-1β expression and increased SQOR expression.

## Discussion

### Fluctuating high temperatures induce a stress response in zebrafish embryos

Communication between conspecifics can potentiate the response of a group to stressors (Giacomini et al., 2015), consequently the combined effects of abiotic stressors and any feedback loops they induce are of great importance in the context of global change. For instance, by the end of the century not only temperature but also chemical cues associated with a warming aquatic environment could be altered (Chivers et al., 2013; Lienart et al., 2016; Roggatz et al., 2019), and warming is known to affect predator-prey communication via infochemicals (J. Xia et al., 2020). The overarching aim of this work was to investigate whether zebrafish embryos can propagate aspects of their response to fluctuating heat stress to naive receiver embryos through positive feedback loops.

First, we investigated the effects of fluctuating thermal stress on zebrafish embryos. Our results showed that heat stressed embryos grew faster than control embryos, which is consistent with previous reports (Long et al., 2012; Schnurr et al., 2014). This may in turn favour premature hatching of smaller larvae (Cingi et al., 2010; Schmidt & Starck, 2010). Stressed embryos were less active than control embryos incubated for 24 hours but were hyperactive compared to controls developed to the same stage of prim-6 without stress treatments. Our results support previous observations that heat stress triggers higher behavioural activity in zebrafish early stages (Gau et al., 2013; Yokogawa et al., 2014). Such behavioural alterations may be explained by (i) energy trade-offs between behaviour, growth, and the metabolic costs of stress response (Barton & Iwama, 1991; Simčič et al., 2015), as well as (ii) temperature-dependent molecular changes in gene expression, epigenetic gene regulation, or post-translational modification related to behaviour, potentially involving circadian clock and neurodevelopmental genes (Colson et al., 2019; López-Olmeda & Sánchez-Vázquez, 2011).

At the gene expression level, we found that IL-1β and SQOR remained unchanged in thermally-stressed embryos. This contrasts with previous findings of SQOR upregulation in response to thermal and hypoxic stresses (Guo et al., 2016; Long et al., 2012, 2015; J. H. Xia et al., 2018) and of heat-induced increased IL-1β levels in adult black rockfish (*Sebastes schlegelii*) (Lyu et al., 2018) and zebrafish embryos (Icoglu Aksakal & Ciltas, 2018). Higher temperature stress usually triggers an upregulation of SOD1 (Liu et al., 2018; Mahanty et al., 2016). In contrast, fluctuating thermal stress in this study reduced SOD1 expression compared to the control, measured almost eight hours after the cessation of thermal fluctuations. These inverted gene expression patterns under fluctuating, as compared to constant thermal stress, might be related to energetic depletion as a result of the thermal cycles (Schaefer & Ryan, 2006). Repetition of heat stress also downregulated heat shock proteins in lake whitefish (*Coregonus clupeaformis*) embryos (Whitehouse et al., 2017). Nevertheless, it should be acknowledged that the differences in developmental rate observed under fluctuating heat stress may superimpose confounding effects to the effects of the heat stimulus, as gene expression varies with ontogeny during zebrafish embryogenesis (Mathavan et al., 2005). Nonetheless, it is likely that fluctuating temperatures experienced through early development of zebrafish embryos cause lasting developmental and behavioural changes, similar to those already shown to occur in vertebrate ectotherms after static increased temperatures.

### Thermal stress induces positive stress feedback loops in naive receiver embryos

It is well-known that animals can chemically communicate a state of distress to others, although predation stress has traditionally received the most attention (Barcellos et al., 2014; Jordão & Volpato, 2000). Here, we found that fluctuating thermal stress negatively affects naive conspecifics, with a similar directionality of effects than the thermal stress itself, which can be described as a positive feedback loop.

First, embryos in our experiment grew faster when subjected to stress medium obtained from previously heat-stressed embryos, reaching similar stages to those of heat-stressed embryos. Such “catch-up” synchronous growth has been shown in egg-clustered embryos of several species and indicates the presence of embryo-embryo communication. Usually this communication serves to maximise energy costs against potential threats and to potentially facilitate group emergence (Aubret et al., 2016; Colbert et al., 2010; McGlashan et al., 2012).

Second, stress medium triggered behavioural hyperactivity compared to control embryos reaching an equivalent stage (prim-6). These results are in agreement with higher activities in rainbow trout embryos facing alarm cues (Poisson et al., 2017) but depart from lower behaviour activities in 24 hpf zebrafish embryos exposed to conspecific alarm cues (Lucon-Xiccato et al., 2020), which suggest that the response depends on the type of the cue and the tested model. In our experiment, these behavioural responses were measured, for the first time, as a response to temperature stress-elicited cues. Behavioural alteration was also observed in adult zebrafish in response to low pH and fasting stress-induced metabolites (Abreu et al., 2016), and adult marine invertebrates experiencing metabolites from low pH-stressed conspecifics and heterospecifics (Feugere et al, in review).

Third, stress medium originating from thermally-stressed embryos induced changes in gene expression patterns in naive conspecific receiver embryos. IL-1β was significantly downregulated in stress medium treatments. The stress medium-mediated inhibition of IL-1β in our experiment suggests that one of its inhibitor stress hormones, such as cortisol, adrenaline and the adrenocorticotropic hormone (Castillo et al., 2009; Castro et al., 2011), or HSF1 (Cahill et al., 1996) could be upregulated by stress media. Intriguingly, the expression of immune genes may be associated with behavioural changes in zebrafish, since highly responsive fish also have higher IL-1β expression (Kirsten et al., 2018). This indicates a functional link between the concomitant decreases of IL-1β and lower activity of stress medium-treated embryos compared to 24 hrs-incubated medium control embryos.

On the other hand, SQOR expression was significantly upregulated by stress medium. SQOR has been little studied in an environmental stress response context thus far, but emerged as a novel candidate marker from recent transcriptomics studies of thermal and oxidative stress (Guo et al., 2016; Long et al., 2012, 2015; Wollenberg Valero et al., 2019; J. H. Xia et al., 2018). SQOR is involved in the metabolism of hydrogen sulfide (H_2_S), concentrations of which are toxic at supraphysiological levels, by tuning its neuromodulatory and biological roles (Augustyn et al., 2017; Chao et al., 2012; Horsman & Miller, 2016; Jackson et al., 2012; Rose et al., 2017). Interestingly, SQOR and H_2_S may be involved in the response to oxidative stress through increasing glutathione levels (Kimura et al., 2010; Yonezawa et al., 2007) and by mediating the antioxidant effects of CoQ10 (Kleiner et al., 2018). There is evidence that SQOR maintains ATP production (Quinzii et al., 2017) and has been proposed as a growth-related candidate gene (Zhuang et al., 2020). Kleiner and colleagues (Kleiner et al., 2018) found that an increase of SQOR may prevent oxidative stress by facilitating the antioxidant effects of CoQ10. Reversely, a downregulation of SQOR may reflect deficiency of its coenzyme CoQ10, which in turn alters the sulfide metabolism leading to accumulated H_2_S levels and depletion of glutathione, that may cause oxidative damages (Luna-Sánchez et al., 2017; Quinzii et al., 2017; Ziosi et al., 2017). Therefore, the upregulation of SQOR under stress medium treatment could have multiple functions, from metabolising toxic levels of H_2_S to restoring both ATP and GSH levels in response to stress communication from a conspecific. Altogether, our results indicate impaired immune and antioxidant responses in embryos exposed to propagated thermal stress.

### Stress medium and control medium do not induce similar feedback mechanisms

Embryos exposed to the control medium developed slower than control embryos from E3 medium, showing opposite directionality to all other treatments. This opposite directionality was mirrored by the expression patterns of the three investigated genes. The behavioural response to stress medium was also significantly less pronounced than that to control medium, corroborating previous studies where metabolites from undisturbed versus stressed donors induced different responses in several species (Bairos-Novak et al., 2017; Bett et al., 2016). However, activity was much higher in control embryos from E3 medium than in embryos incubated in control medium. This finding may indicate that behavioural activity of zebrafish embryos is tightly controlled by the nature of their chemical environment (vs. relaxed in the absence of any cues), lending additional support to chemical communication as a global change-relevant parameter.

Lastly, the similarity in development and behavioural values in combined thermal stress plus stress medium treatments together with a larger effect size indicate that thermal stress and stress medium may excite similar molecular pathways regulating growth rate and behavioural activity, but that these pathways are saturated by single stressors, and cannot be further altered in the combined treatment. Conversely, our gene expression analysis revealed a difference in molecular responses between the two independent factors, thermal stress and stress medium. To better understand these contrasting synergistic vs. independent effects of thermal stress and stress medium, gene expression could be compared at global scale in future work.

### Conclusion

To conclude, our study indicates that thermally-stressed zebrafish embryos induce a stress response in naive conspecifics that have not been exposed to thermal stress, from molecular to phenotypic level. This extends the concept of stress feedback loops and chemical communication of stress to fluctuating heat stress in a vulnerable life stage of fish. This is important because such stress feedback loops may be widespread and have implications on tipping points of ecosystems facing ongoing and future global change. We suggest further investigation into the identification of the nature of metabolites contained within stress medium, their molecular consequences at an individual level, as well as longer-term consequences for populations and ecosystems.

## Acknowledgements

The authors acknowledge Alan Smith and Sonia Jennings for providing fish care, Emma Chapman, Robert Donnelly for support in the laboratory, Adam Bates and Paulo Saldanha (University of Hull, UK) for support in discussing the experiments, and all members of the MolStressH2O research cluster (University of Hull, UK) for valuable discussions and peer review. For help with initial zebrafish method development we thank Antoinette Destefano, Amber Pinnock, Tiana Weeks, and Tia Rusciano (Bethune-Cookman University, USA) and Tyrese Taylor (Indiana University, USA). Funding was provided by the University of Hull to LF, VS, QRB, PBA and KWV and the Royal Society (RGS\R2\180033) to KWV.

## Ethics approval

All experiments were approved by the University of Hull Ethics committee under the approval U144b.

## Consent to participate

All authors consented to participation.

## Consent for publication

All authors consented to publication.

## Author contribution

KWV conceived the study. KWV and PBA designed the experiments. LF, QRB, and VS collected the data and contributed to the statistical analysis. LF wrote the manuscript draft with PBA and KWV. All authors contributed to the final manuscript.

