## Supplementary Information for "Thermal stress induces positive phenotypic and molecular feedback loops in zebrafish embryos"

### Supplementary Information to the Manuscript '*Thermal stress induces positive phenotypic and molecular feedback loops in zebrafish embryos*'

Lauric Feugere[1], Victoria F. Scott[1], Quentin Rodriguez-Barucg[2], Pedro Beltran-Alvarez[2] & Katharina C. Wollenberg Valero[1]*

[1]Department of Biological and Marine Sciences, University of Hull, Cottingham Road, Kingston-upon-Hull HU6 7RX, United Kingdom.
[2]Department of Biomedical Sciences, University of Hull, Cottingham Road, Kingston-upon-Hull HU6 7RX, United Kingdom.

#### Supplementary Methods

##### Animals and breeding

Adult zebrafish (*Danio rerio*) were kept in optimal conditions (controlled temperature: 28 ± 1°C, 12 hrs light/12 hrs dark cycle, pH = 7.6, salinity = 1) at the Aquarium facility of the University of Hull. Adult zebrafish were fed twice daily an alternate diet of bloodworm and *Daphnia* plus vitamins. Plastic trays half-filled with marbles and plastic plants were left overnight in the fish tank for breeding (Nasiadka & Clark, 2012). Breeders were randomly selected from the colony and allowed to rest for at least two days between breeding events. In the morning, eggs were collected and transferred into the laboratory. Experimental treatments were conducted on embryos collected at different times throughout the year 2019. All experiments were approved by the Ethics committee of the University of Hull (Ethics form U144b). Just after collection, eggs were rinsed in tap water and immediately placed in 1X E3 medium (Cold Spring Harbor Laboratory Press, 2011) containing 1% methylene blue. Embryos were observed under a stereomicroscope (ZEISS SteREO Discovery.V8, Carl Zeiss) mounted with an Axiocam 105 Color camera (Carl Zeiss) and coupled with the ZEN lite software (Carl Zeiss).

##### Total RNA extraction

After experimental treatments, fertilised embryos were individually transferred from their holding exposure well to a nuclease-free microcentrifuge tube and medium was removed. Samples were prepared from pools of 60 embryos, to obtain an adequate amount of tissues for kit-based RNA extraction. Embryos were humanely euthanized by snap freezing at -80°C and stored until further processing. RNA was extracted from several samples per treatment and the best 3 samples were kept for further analysis. Total RNA was extracted using the High Pure RNA Isolation Kit (cat. n° 12033674001, Sigma-Aldrich Company Ltd., Missouri, USA) following the manufacturer's recommendations albeit small adjustments were made for yield optimisation, to the following steps: (i) 400 µL of lysis/binding buffer (in two steps of 100 µL and 300 µL with 15 min incubation in ice for optimal tissue disruption) were added to 1.5 mL tubes and embryos were manually homogenised for 30-45 s using plastic pestles (pre-cleaned with RNAse-away and 75% ethanol). (ii) Total RNA was eluted in 50 µL Elution Buffer and centrifuged 1 min at 8,000 rpm. Eluted RNA was passed another time through the column and the centrifugation repeated to enhance extraction yields. For each reaction, 11 µL were aliquoted for triple-assessment of RNA quality whilst the remaining volume was immediately stored at -80°C until further processing to prevent RNA degradation. RNA integrity was assessed by observing clear 28S and 18S bands in 1% agarose gels. Total RNA was quantified using the Qubit® fluorometer and Qubit RNA BR Assay Kit (Invitrogen). RNA purity was assessed using the NanoDrop 1000 spectrophotometer (Thermo Scientific). Ratios ranged within (2.03-2.30) and (1.98-2.22) for 260/280 and 260/230 ratios, respectively. Total RNA was deemed suitable for further processing.

**Complementary DNA synthesis**

After sample randomisation, mRNA samples were reverse transcribed in 20 uL of reaction for a final concentration of 16 ng/uL using the Superscript II ™ Reverse Transcriptase kit (Invitrogen, Life Technologies Ltd.). For each reaction, 320 ng of RNA were combined with 1 µL of oligodT (Nanoscript Precision™ OligoDT primer mix, Primer Design), 1 µL of nucleotides (dNTP mix, 10 mM each), and molecular grade water up to a total volume of 13 µL. The preparation was incubated 5 min at 65°C and immediately chilled in ice. Four microliters of 5X First Strand Buffer and 2 µL of 0.1 M DTT (Invitrogen, Life Technologies Ltd.) were added to each reaction. After incubation at 42°C for 2 min, 1 µL Superscript II ™ Reverse Transcriptase (Invitrogen, Life Technologies Ltd.) was added. No Reverse Transcription (NRT) reactions were prepared by replacing the enzyme by molecular grade water. cDNA was then synthesised for 50 min at 42°C and the reaction terminated by 15 min at 70°C. The success of the cDNA synthesis was checked (1) by 1% agarose gel electrophoresis and (2) by conventional PCR with previously optimised custom primers for elongation factor alpha 1 (GenBank: NM_131263, forward primer: 5'-GATGCACCACGAGTCTCTGA-3'; reverse primer: 5'-TGATGACCTGAGCGTTGAAG-3') and Cu/Zn-superoxide dismutase (GenBank: Y12236, forward primer: 5'-CGTCTGGCTTGTGGAGTGAT-3'; reverse primer: 5'-TAATGTCAGCGGGCTAGTGC-3'). The PCR program consisted of 5 min at 95°C followed by 40 cycles of 30 s at 95°C, 30 s at 62°C, and 30 s at 72°C, 1 min at 60°C and a final elongation of 5 min at 72°C. PCR reactions were loaded in a 2% agarose gel in 1X TBE alongside 100p DNA ladder (New England Biolabs). The presence of one sharp band in +RT and the absence of amplicon in NRT samples validated the cDNA integrity. Sharp bands were observed when loading cDNA from 0.16 ng to 32 ng in 20 µL of PCR reaction, which justified dilution of cDNA samples for qPCR. Four-fold diluted cDNA samples (4 ng/µL) were then prepared in molecular grade water and stored at -20°C until used in qPCR.

**Quantitative Real-Time PCR**

The expression of three genes of interest (GOIs) (sulfide:quinone oxidoreductase, SQOR, TaqMan® assay ID: Dr03122210_m1; superoxide dismutase 1 soluble, SOD1, TaqMan® assay ID: Dr03074067_m1; interleukin 1, beta, il1b, TaqMan® assay ID: Dr03114368_m1, here referred to as IL-1β) was measured. Two reference genes (eukaryotic translation elongation factor 1 alpha 1, like 1, eef1a1l1 - here referred to as ef1-ɑ, Dr03432748_m1; and actin, beta 1, actb - here referred to as β-actin, Dr03432610_m1) were chosen for normalisation. The compatibility of the universal probe kit 2X qPCR Bio Probe Hi-ROX (PCRBiosystems) with each TaqMan probe was assessed after successful amplification in conventional PCR. For this, half reactions were prepared with 2 µL of diluted cDNA samples (4 ng/µL), 0.5 µL of 20✕ TaqMan® Gene Expression Assay, and 5 µL of 2X qPCR Bio Probe Hi-ROX. The PCR program was as follows: 5 min at 50°C followed by 40 cycles of 15 s at 95°C and 1 min at 60°C. PCR samples were loaded in 2% agarose gels in 1X TBE. Sharp bands were observed for diluted cDNA samples and NRT samples did not result in amplification. Gene expression was subsequently measured in quantitative PCR with methods based on TaqMan® Gene Expression Assays (Thermofisher, available at http://tools.thermofisher.com/content/sfs/manuals/cms_041280.pdf). Each 20 µL qPCR reaction contained 16 µL of a master mix made of 10 µL of 2X qPCR Bio Probe Hi-ROX (PCRBiosystems), 1 µL of 20✕ TaqMan® Gene Expression Assay, and 5 µL of molecular grade water. Each qPCR reaction was completed with 4 µL of cDNA template at 4 ng/µL. No template Control (NTC) and NRT reactions were added and validated the procedure as they were not expressed or at > 5 Ct after the lowest sample. Standard runs (2 hrs) of qPCR consisted of 2 min at 50°C, 10 min at 95°C, followed by 40 cycles of 15 s at 95°C and 1 min at 60°C. Gene expression was analysed using the StepOnePlus™ Real-Time PCR System coupled with the StepOne™ software v2.3 (Applied Biosystems™). Sample maximisation strategy (as many samples as possible for one gene measured in the same plate) was used to limit run to run variations (Hellemans et al., 2007). Reference gene stability was verified using RefFinder (available online at https://www.heartcure.com.au/reffinder/, accessed January 2020, (Xie et al., 2012)). GeNorm (Vandesompele et al., 2002), NormFinder (Andersen et al., 2004), and BestKeeper (Pfaffl et al., 2004) algorithms were used to assess the suitability of reference genes. GeNorm expression stability value M was below 0.5 and 1.5 (M = 0.164 for ef1-ɑ and geometric mean, M = 0.239 for β-actin) which indicated stable reference genes (D'haene & Hellemans, 2010; Sarker et al., 2018). Standard deviations of cycle threshold were around 1 (SD Ct ef1-ɑ = 1.011, SD Ct β-actin = 1.098, SD Ct geometric mean = 1.042) which was deemed stable (Pfaffl et al., 2004). The geometric mean of both reference genes decreased the stability value in normFinder (ef1-α: 0.243, β-actin: 0.264, geometric mean: 0.082) and was used to normalise the gene expression of GOIs. Cycle thresholds of samples that were run on different plates were expressed as calibrated Ct (CCt) values (Vermeulen et al., 2009) relative to one or two intercalibrators (IRC). The relative gene expression was expressed as $Log_2\ 2^{-\Delta\Delta CT}$ where $\Delta\Delta CT = (Ct\ GOI - Ct\ ref)_{treatment} - (Ct\ GOI - Ct\ ref)_{Control}$ (Livak & Schmittgen, 2001).

**Data analysis**

Additional to temperature and stress medium as the main predictors, the effects of confounding factors were assessed. Because experiments were repeated on different days, median initial stages slightly differed across batches with CM treatments starting in average at 2.75 hpf (512-cell) and thus were somewhat older than in C and SM conditions that started at 2 hpf in average (64-cell). However, SM, TS, and TS+SM treatments started at similar initial stages compared to control embryos. The effects of batches' median initial stages on behavioural responses were analysed using Kruskal-Wallis tests.

#### Supplementary Results

**Effects of thermal stress, stress medium, and confounding factors on growth and activity**

As a consequence of faster growth, thermally-stressed embryos (H = 71.096, P < 0.0001) and the interaction of thermal stress and stress medium (H = 6.074, P = 0.0137) led to significantly older embryonic stages (Figure S1a, Table S3). Embryos stressed by SM (W = 611.5, P = 0.0056), TS (W = 766.5, P < 0.0001), and TS+SM (W = 526.5, P < 0.0001) treatments were at a median stage of 25 hpf (prim-6) whereas control embryos were 22-hpf old (26-somite). Embryos in CM were also 22-hpf, that is similar to C but significantly younger than SM (W = 699, P = 0.0015). (Kimmel et al., 1995) ranged segmentation and pharyngula embryonic periods as ]10-22 hpf] and [24-48 hpf[, respectively. TS (Z = -7.035, P < 0.0001), SM (Z = -3.255, P = 0.0011), and the interaction of TS + SM (Z = 2.897, P = 0.0038) significantly shifted the timing of entrance into the pharyngula period (Figure S1b, Table S4). Most of stressed embryos were in pharyngula period after treatment with SM (Z = -3.255, P = 0.0034), TS (Z = -7.035, P < 0.0001), and TS+SM (Z = -5.804, P < 0.0001) against only a quarter in C. The percentage of embryos in the pharyngula period in CM did not significantly vary compared to control but was significantly lower than in SM (Z = 3.456, P = 0.0005). Final stages ($\chi^2$ = 104.49, Spearman's ρ = 0.25, both statistics with P < 0.0001) and period (Z = -2.5211, P = 0.0117) were significantly dependent on median initial stages which indicates that stage shifts due to treatment scale across different initial stages.

In addition to the effects of both factors (thermal stress and stress medium) mentioned in the main result section, burst activity percentages significantly decreased with initial stages (Spearman's ρ = -0.33, P < 0.001) across all treatments. However, burst activity percentages did not correlate with final stages (in hpf, Spearman's ρ = -0.08, P = 0.1627) nor growth indexes (Spearman's ρ =0.04, P = 0.5027, Table S5).

#### Supplementary Figures

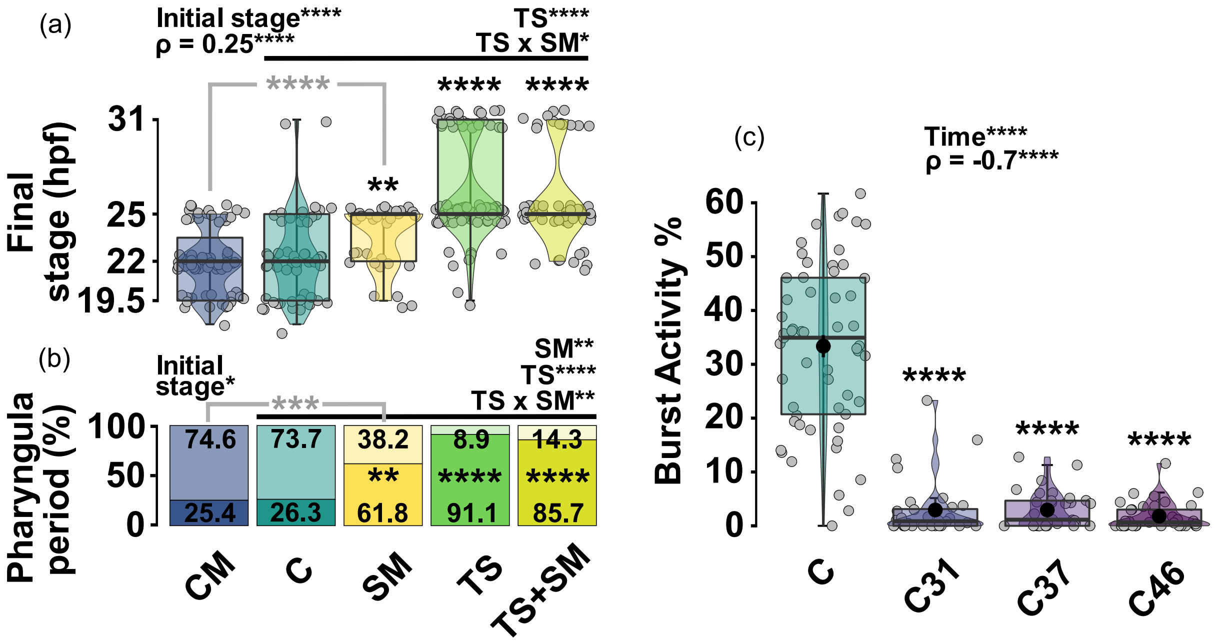


**Figure S1. Thermal stress and stress medium shift the final embryonic stage, and activity decreases throughout pre-hatching development in control embryos.** (a) Final stage in hour post fertilisation (hpf). (b) Percentage of embryos reaching the pharyngula stage. Final stage was binary coded as segmentation (18 to 22 hpf, top light areas) and pharyngula (25 to 42 hpf, bottom dark areas shown as percentages) periods. (c) Burst activity percentage of pre-hatching embryos at different times of incubation in control temperature of 27°C. Boxes represent median, 25%-75% quartiles, and whiskers are minimum and maximum values within 1.5 IQR (interquartile range). Jittered raw data given as grey circles. Black dots represent the mean ± SE values of each treatment. Left panel: CM: control medium ($n$ = 67), C: control ($n$ = 57), SM: stress medium ($n$ = 34), TS: thermal stress ($n$ = 90), TS+SM: thermal stress + stress medium ($n$ = 56). Effects of thermal stress (TS) and stress medium (SM) predictors were tested with a non-parametric factorial two way Scheirer-Ray-Hare or a generalised linear models for logistic distribution, when final stages were expressed as hpf or binary period, respectively (significant predictors shown on top right corners above horizontal black lines). Significance of pairwise post-hoc tests is likewise indicated in asterisks, for treatments against C (bold black), between SM and CM (light grey). Stages were determined with reference to (Kimmel et al., 1995). Right panel: control embryos were incubated starting approx. 2.75 hrs post fertilisation for 24 hrs (C, $n$ = 57), 31 hrs (C31, $n$ = 32), 37 hrs (C37, $n$ = 32), and 46 hrs (C46, $n$ = 37). Significance of pairwise post-hoc tests is indicated in asterisks, between control C and C31, C37, or C46. In both panels, effects of initial stages were analysed with Kruskal-Wallis' tests and Spearman's rho correlation test. *: $P \leq 0.05$, **: $P \leq 0.01$, ***: $P \leq 0.001$, ****: $P \leq 0.0001$.

#### Supplementary Tables

**Table S1. Details of experimental treatments.** Zebrafish embryos (*Danio rerio*) in treatments for 24 hours starting approx. 2.75 hrs post fertilisation**.** The design aimed to investigate the effects of fluctuating thermal stress and stress medium excreted by stressed embryos. In addition, control embryos were incubated at 27°C for different incubation times starting approx. 2.75 hrs post fertilisation. Hpf: hours post fertilisation. The final behavioural dataset contained 405 observations (C: $n$ = 57; CM: $n$ = 67; TS: $n$ = 90; SM: $n$ = 34; TS+SM: $n$ = 56; C-31hrs: $n$ = 32; C-37hrs: $n$ = 32; C-46hrs: $n$ = 37). Gene expression analysis was conducted on CM, C, SM, TS, TS+SM with 3 samples containing 60 embryos per treatment.

| Treatments | Detail |
|---|---|
| Control (C) | Incubation at 27°C starting 3.3 hpf for 24 hrs (from 11 am at day 1 to 11 am at day 2) in the dark. Incubation in fresh 1X E3 medium. Endpoints: qPCR, behaviour, growth. |
| Control Medium (CM) | Incubation at 27°C starting 3.3 hpf for 24 hrs (from 11 am at day 1 to 11 am at day 2) in the dark. Incubation in reused 1X E3 medium, in which an embryo from the C treatment was exposed beforehand and containing putative control metabolites. Endpoints: qPCR, behaviour, and growth. |
| Thermal Stress (TS) | Incubation in thermal stress treatment starting 3.3 hpf for 24 hrs (from 11 am at day 1 to 11 am at day 2) in the dark. Thermal stress spanned 16.25 hrs, divided into thirteen 75-min series of temperature fluctuations between 27, 29, 32, 29, and 27°C, with each temperature step being maintained for 15 min. Incubation in fresh 1X E3 medium. Endpoints: qPCR, behaviour, growth. |
| Stress Medium (SM) | Incubation at 27°C starting 3.3 hpf for 24 hrs (from 11 am at day 1 to 11 am at day 2) in the dark. Incubation in reused 1X E3 medium, in which an embryo from the TS treatment was exposed beforehand and containing putative stress metabolites. Endpoints: qPCR, behaviour, growth. |
| Thermal Stress + Stress Medium (TS+SM) | Incubation in thermal stress treatment starting 3.3 hpf for 24 hrs (from 11 am at day 1 to 11 am at day 2) in the dark. Incubation in reused 1X E3 medium, in which an embryo from the TS treatment was exposed beforehand and containing putative stress metabolites. Endpoints: qPCR, behaviour, growth. |
| C31, C37, and C46 | Incubation at 27°C from 3.3 hpf to 25 hpf (reached after approx. 31 hrs, prim-6), 31 hpf (reached after approx. 37 hrs, prim-16), or 35-42 hpf (reached after approx. 46 hrs, late pharyngula) in the dark. Exposure in fresh 1X E3 medium. Endpoints: behaviour, growth. |

**Table S2. Effects of thermal stress and stress medium on the growth index of zebrafish embryos (*Danio rerio*).** Embryos were incubated in treatments for 24 hours starting approx. 2.75 hrs post fertilisation (hpf). Growth index was calculated as Δ(final stage - initial stage)/incubation time. C: control ($n$ = 57), SM: stress medium ($n$ = 34), TS: thermal stress ($n$ = 90), TS+SM: thermal stress + stress medium ($n$ = 56), CM: control medium (n = 67). Pairwise Student's tests with Bonferroni adjustments were used to compare all treatments to C and SM to CM. Effect size is computed as Cohen's |d| which can be interpreted according to thresholds given in Sawilowsky (2009). Effect sizes of significant p-values ($P \leq 0.05$) are shown in bold.

| Pairwise comparisons | T statistic | P | \|d\| | Effect size |
|---|---|---|---|---|
| C-SM | -3.6784 | 0.0012 | **0.75** | **medium** |
| C-TS | -7.9874 | 0.0001 | **1.42** | **very large** |
| C-TS+SM | -7.2413 | 0.0001 | **1.36** | **very large** |
| C-CM | 2.3472 | 0.0418 | **0.43** | **small** |
| SM-CM | -6.7215 | 0.0001 | **1.45** | **very large** |

**Table S3. Effects of thermal stress and stress medium on the final stage (hour post fertilisation) of zebrafish embryos (*Danio rerio*).** Embryos were incubated in treatments for 24 hours starting approx. 2.75 hrs post fertilisation. Final stages were expressed as hour post fertilisation according to Kimmel et al. (1995). C: control ($n$ = 57), SM: stress medium ($n$ = 34), TS: thermal stress ($n$ = 90), TS+SM: thermal stress + stress medium ($n$ = 56), CM: control medium (n = 67). A non-parametric two-way Scheirer-Ray-Hare test was used to test the effects of predictors (thermal stress and stress medium) across C, SM, TS, and TS+SM. Pairwise Wilcoxon-Mann-Whitney's tests with Bonferroni adjustments were used to compare all treatments to C and SM to CM. Effect of clutch median initial stages across all treatments were analysed with a Kruskal-Wallis' test whereas correlation with final stage was estimated with a Spearman's test. Effect size is computed as Cohen's |d| which can be interpreted according to thresholds given in Sawilowsky (2009). Effect sizes of significant p-values ($P \leq 0.05$) are shown in bold.

| | **H statistic** | **P** | **\|d\|** | **Effect size** |
|---|---|---|---|---|
| **Model terms** | | | | |
| Stress Medium | 1.221 | 0.2693 | 0.10 | very small |
| Thermal Stress | 71.096 | 0.0001 | **1.23** | **very large** |
| Thermal Stress x Stress Medium | 6.074 | 0.0137 | **1.25** | **very large** |
| **Pairwise comparisons** | **W statistic** | **P** | **\|d\|** | **Effect size** |
| C-SM | 611.5 | 0.0056 | **0.56** | **medium** |
| C-TS | 766.5 | 0.0001 | **1.42** | **very large** |
| C-TS+SM | 526.5 | 0.0001 | **1.34** | **very large** |
| C-CM | 1893 | 1.000 | 0.07 | very small |
| SM-CM | 699 | 0.0015 | **0.75** | **medium** |
| **Covariate** | **Kruskal-Wallis' test terms** | | **Spearman's test terms** | |
| Initial stage | $X^2 = 104.59$, $P < 0.0001$ | | $\rho = 0.25$, $P < 0.0001$ | |

**Table S4. Effects of thermal stress and stress medium on the final stage period of zebrafish embryos (*Danio rerio*).** Embryos were incubated in treatments for 24 hours starting approx. 2.75 hrs post fertilisation. Final stages were expressed as binary embryonic periods (segmentation, pharyngula) according to (Kimmel et al., 1995). CM: control medium (n = 67), C: control ($n$ = 57), SM: stress medium ($n$ = 34), TS: thermal stress ($n$ = 90), TS+SM: thermal stress + stress medium ($n$ = 56). A generalised linear model for logistic regression was used to test the effects of (i) thermal stress and medium predictors across C, SM, TS, and TS+SM, and (ii) the effects of embryo clutch median initial stages across all treatments. Pairwise tests with Bonferroni adjustments were computed using the *emmeans* r package to compare all treatments to C and SM to CM. Effect size is computed as Cohen's |d| which can be interpreted according to thresholds given in Sawilowsky (2009). Significant p-values ($P \leq 0.05$) are shown in bold.

| **Model terms** | **Estimate** | **SE** | **Z statistic** | **P** |
|---|---|---|---|---|
| Stress Medium | -1.5092 | 0.4637 | -3.255 | **0.0011** |
| Thermal Stress | -3.3569 | 0.4771 | -7.035 | **0.0001** |
| Thermal Stress x Stress Medium | 2.0447 | 0.7057 | 2.897 | **0.0038** |
| **Pairwise comparisons** | | | | |
| C-SM | -1.51 | 0.464 | -3.255 | **0.0034** |
| C-TS | -3.36 | 0.477 | -7.035 | **0.0001** |
| C-TS+SM | -2.82 | 0.486 | -5.804 | **0.0001** |
| C-CM | -0.05 | 0.411 | -0.120 | 1.000 |
| SM-CM | 1.56 | 0.451 | 3.456 | **0.001** |
| **Covariate** | | | | |
| Initial stage | -0.4056 | 0.1609 | -2.5211 | **0.0117** |

**Table S5. Effects of thermal stress and stress medium on the burst activity percentage of zebrafish embryos (*Danio rerio*).** Embryos were incubated in treatments for 24 hours starting approx. 2.75 hrs post fertilisation. Burst activity percentages were measured from videos using Danioscope. CM: control medium (n = 67), C: control (*n* = 57), SM: stress medium (*n* = 34), TS: thermal stress (*n* = 90), TS+SM: thermal stress + stress medium (*n* = 56). Pairwise Wilcoxon-Mann-Whitney's tests with Bonferroni adjustments were used to compare all treatments to C and SM to CM. Spearman's correlation tests were used to estimate the confounding effects of covariates (clutch median initial stage, individual final stage expressed as hours post fertilisation, hpf, or growth index) on burst activity percentages. Effect size is computed as Cohen's |d| which can be interpreted according to thresholds given in Sawilowsky (2009). Effect sizes of significant p-values (P ≤ 0.05) are shown in bold.

| **Pairwise comparisons** | **W statistic** | **P** | **\|d\|** | **Effect size** |
|---|---|---|---|---|
| C-SM | 1387 | 0.0018 | **0.81** | **large** |
| C-TS | 3548 | 0.0003 | **0.68** | **medium** |
| C-TS+SM | 2455 | 0.0001 | **1.04** | **large** |
| C-CM | 3287 | 0.0001 | **1.64** | **very large** |
| SM-CM | 1745 | 0.0001 | **1.05** | **large** |
| **Covariates** | **ρ statistic** | **P** | - | - |
| Initial stage | -0.33 | 0.0001 | - | - |
| Final stage (in hpf) | -0.08 | 0.1627 | - | - |
| Growth index | 0.04 | 0.5027 | - | - |

**Table S6. Effects of incubation time on the burst activity percentage of control zebrafish embryos (*Danio rerio*).** Embryos were raised at control temperature of 27°C starting approx. 2.75 hrs post fertilisation for 24 hrs (C, *n* = 57), 31 hrs (C31, *n* = 32), 37 hrs (C37, *n* = 32), and 46 hrs (C46, *n* = 37). A non-parametric Kruskal-Wallis test was used to test the effects of incubation times. Pairwise Wilcoxon-Mann-Whitney's tests with Bonferroni adjustments were used to compare all C31, C37, and C46 to C. Effect size is computed as Cohen's |d| which can be interpreted according to thresholds given in Sawilowsky (2009). Effect sizes of significant p-values (P ≤ 0.05) are shown in bold.

| **Model term** | **$X^2$ statistic** | **P** | **-** | **-** |
|---|---|---|---|---|
| Incubation time | 99.639 | 0.0001 | - | - |
| **Pairwise comparisons** | **W statistic** | **P** | **\|d\|** | **Effect size** |
| C-C31 | 1762 | 0.0001 | **2.37** | **huge** |
| C-C37 | 1775 | 0.0001 | **2.41** | **huge** |
| C-C46 | 2065 | 0.0001 | **2.59** | **huge** |
| **Correlation test** | **ρ statistic** | **P** | **-** | **-** |
| Incubation time | -0.70 | 0.0001 | - | - |

**Table S7. Effects of thermal stress and stress medium on the burst activity percentage of zebrafish embryos (*Danio rerio*) at the median stage of prim-6.** Comparison of the burst activity percentage of zebrafish embryos reaching the median stage of prim-6 in control condition (C31) and stress treatments (SM, TS, TS+SM) after 31 hrs and 24 hrs of incubation, respectively. C31: control (*n* = 32), SM: stress medium (*n* = 34), TS: thermal stress (*n* = 90), TS+SM: thermal stress + stress medium (*n* = 56). Pairwise Wilcoxon-Mann-Whitney's tests with Bonferroni adjustments were used to compare all treatments to C31. Effect size is computed as Cohen's |d| which can be interpreted according to thresholds given in Sawilowsky (2009). Effect sizes of significant p-values (P ≤ 0.05) are shown in bold.

| **Pairwise comparisons** | **W statistic** | **P** | **\|d\|** | **Effect size** |
|---|---|---|---|---|
| C31-SM | 37 | 0.0001 | **2.63** | **huge** |
| C31-TS | 146 | 0.0001 | **1.64** | **very large** |
| C31-TS+SM | 96 | 0.0001 | **1.75** | **very large** |

**Table S8. Effect of thermal stress and thermal stress medium on the gene expression of zebrafish embryos (*Danio rerio*) exposed during 24 hours.** CM: control medium (n = 67), C: control (*n* = 57), SM: stress medium (*n* = 34), TS: thermal stress (*n* = 90), TS+SM: thermal stress + stress medium (*n* = 56). Pairwise tests of treatments were computed via moderated t-tests (t-statistic), corrected for multiple testing with *lmfit* and *eBayes* in the *limma* R package. Treatments (SM, TS, TS+SM) were compared to control C. Additionally, CM was compared to C and SM. Sample size was n = 3 biological replicates (each containing 60 embryos) per treatment. Effect size is computed as Cohen's |d| which can be interpreted according to thresholds given in Sawilowsky (2009). Effect sizes of significant p-values ($P \leq 0.05$) are shown in bold.

| Pairwise comparisons | | t | B | P | \|d\| | Effect size |
|---|---|---|---|---|---|---|
| **IL-1β** | | | | | | |
| | C-SM | 1.60 | -4.50 | 0.393 | 1.24 | very large |
| | C-TS | 0.75 | -5.63 | 1.00 | 0.82 | large |
| | C-TS+SM | 2.37 | -3.48 | 0.097 | 1.51 | very large |
| | C-CM | 1.02 | -5.77 | 0.639 | 0.89 | large |
| | SM-CM | 2.62 | -3.068 | 0.034 | **1.82** | **very large** |
| **SOD1** | | | | | | |
| | C-SM | 1.59 | -4.51 | 0.401 | 1.03 | large |
| | C-TS | 2.76 | -2.93 | 0.045 | **1.78** | **very large** |
| | C-TS+SM | 1.69 | -4.45 | 0.336 | 1.03 | large |
| | C-CM | -3.78 | -0.992 | 0.003 | **2.20** | **huge** |
| | SM-CM | -2.64 | -3.05 | 0.034 | **1.94** | **very large** |
| **SQOR** | | | | | | |
| | C-SM | -2.22 | -3.67 | 0.13 | 2.09 | huge |
| | C-TS | -0.19 | -5.89 | 1.00 | 0.19 | very small |
| | C-TS+SM | -2.36 | -3.49 | 0.098 | 3.22 | huge |
| | CM-C | -0.62 | -6.09 | 1.00 | 0.77 | medium |
| | CM-SM | -2.20 | -3.80 | 0.08 | 2.34 | huge |
